# A Computational Model of Perceptual and Mnemonic Deficits in Medial Temporal Lobe Amnesia

**DOI:** 10.1101/082628

**Authors:** Patrick S. Sadil, Rosemary A. Cowell

## Abstract

Damage to the Medial Temporal Lobe (MTL) has long been known to impair declarative memory and recent evidence suggests that it also impairs visual perception. A theory termed the representational-hierarchical account explains such impairments by assuming that MTL stores conjunctive representations of items and events, and that individuals with MTL damage must rely upon representations of simple visual features in posterior visual cortex, which are inadequate to support memory and perception under certain circumstances. One recent study of visual discrimination behavior revealed a surprising anti-perceptual learning effect in MTL-damaged individuals: with exposure to a set of visual stimuli, discrimination performance worsened rather than improved (Barense et al., 2012). We extend the representational-hierarchical account to explain this paradox by assuming that difficult visual discriminations are performed by comparing the relative ‘representational tunedness’ – or familiarity – of the to-be-discriminated items. Exposure to a set of highly similar stimuli entails repeated presentation of simple visual features, eventually rendering all feature representations maximally, and thus equally, familiar ― hence, they are inutile for solving the task. Discrimination performance in patients with MTL lesions is therefore impaired by stimulus exposure. Because the unique conjunctions represented in MTL do not occur repeatedly, healthy individuals are shielded from this perceptual interference. We simulate this mechanism with a neural network previously used to explain recognition memory, thereby providing a model that accounts for both mnemonic and perceptual deficits caused by MTL damage with a unified architecture and mechanism.

## INTRODUCTION

The Medial Temporal Lobe (MTL) has long been associated with declarative memory (Scoville & Milner, 1957; Squire & Zola-Morgan, 1991). Recent studies have implicated MTL structures in other functions, such as high-level perception (e.g., Buckley, Booth, Rolls, & Gaffan, 2001; Lee et al., 2005), decision-making (Wimmer & Shohamy, 2012), statistical learning (Schapiro, Kustner, & Turk-Browne, 2012) and imagination (Maguire, Vargha-Khadem, & Hassabis, 2010; Schacter & Addis, 2007). Thus, while the notion of a declarative memory system in the MTL (Squire and Zola-Morgan, 1991; Squire and Wixted, 2011) has provided critical insight into the organization of memory, it no longer fully explains the role of the MTL in cognition. The challenge, now, is to develop mechanistic theories that explain why MTL structures are implicated in both memory and other cognitive functions.

A theory termed the representational-hierarchical account has been put forward to explain both mnemonic and perceptual deficits caused by damage to different structures within MTL (Bussey & Saksida, 2002; Cowell, Bussey, & Saksida, 2006; Kent, Hvoslef-Eide, Saksida, & Bussey, 2016). This account assumes that the ventral visual stream contains a hierarchical organization of representations that continues into the MTL. Early stages of the pathway (e.g., V1, V2, V4) are assumed to represent simple visual features (e.g., color, orientation), whereas more anterior regions are assumed to bring simple features together into conjunctions of increasing complexity (Fig. 1). The hierarchy culminates in the MTL, where conjunctions correspond to whole objects, scenes, or complex episodic events. The representational-hierarchical account claims that conjunctive representations in the MTL are important whenever a cognitive task – perceptual or mnemonic – employs stimuli with overlapping features, such that individual feature representations provide ambiguous information (Bussey & Saksida, 2002).

**Figure 1.**
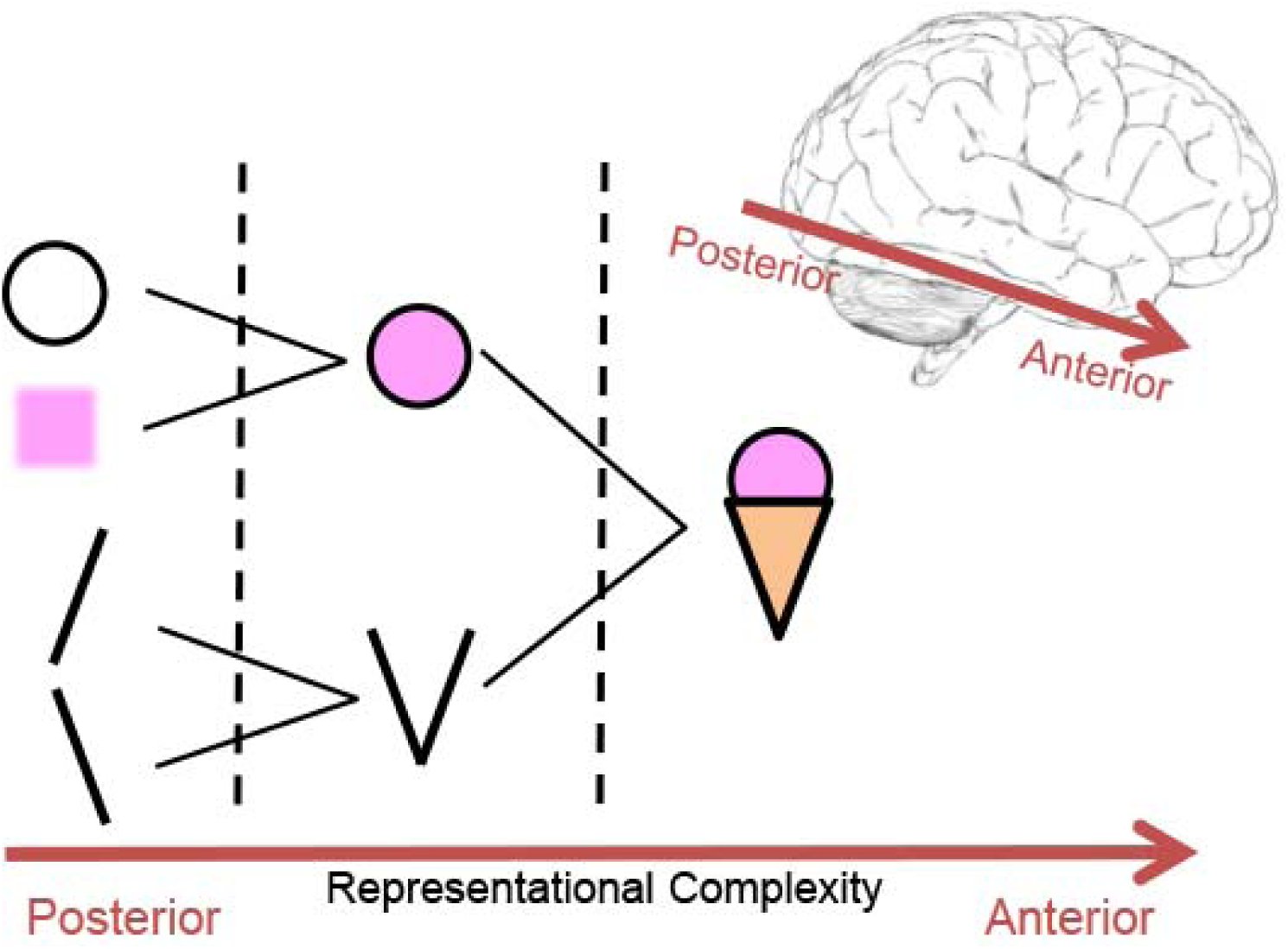
The organization of representations according to the representational-hierarchical account. Simple visual features, such as an oriented line or a color, are represented in early visual regions. Features are combined into increasingly complex conjunctions going from posterior to anterior regions. The MTL lies at the apex of the hierarchy, where conjunctions correspond to whole objects (in perirhinal cortex) or to spatial scenes or episodic events comprising multiple objects and their context (in hippocampus)

### Paradoxical Finding of an Exposure-induced Deficit in Visual Discrimination

A recent study by Barense et al. (2012) documented a new and puzzling way in which MTL lesions impair visual discrimination. MTL amnesics and healthy controls were asked to judge whether pairs of simultaneously presented abstract stimuli were the same or different (Fig. 2a). Stimuli were defined by three features: outer shape, inner shape, and fill pattern. To ensure that participants could not perform the task based on pixel-wise differences between stimuli, the discriminanda were rotated by a random angle between 15° and 165°. The task required participants to declare a mismatch if any of the three features differed across the pair, or a match if the two stimuli were identical. In the High Ambiguity condition, each pair of stimuli in a mismatch trial shared two out of three features, whereas in the Low Ambiguity condition, items in a mismatching pair shared no features. Amnesic patients were unimpaired at discriminating Low Ambiguity objects, but for High Ambiguity discriminations the performance of MTL patients deteriorated in the second half of trials (Fig. 2b).

**Figure 2.**
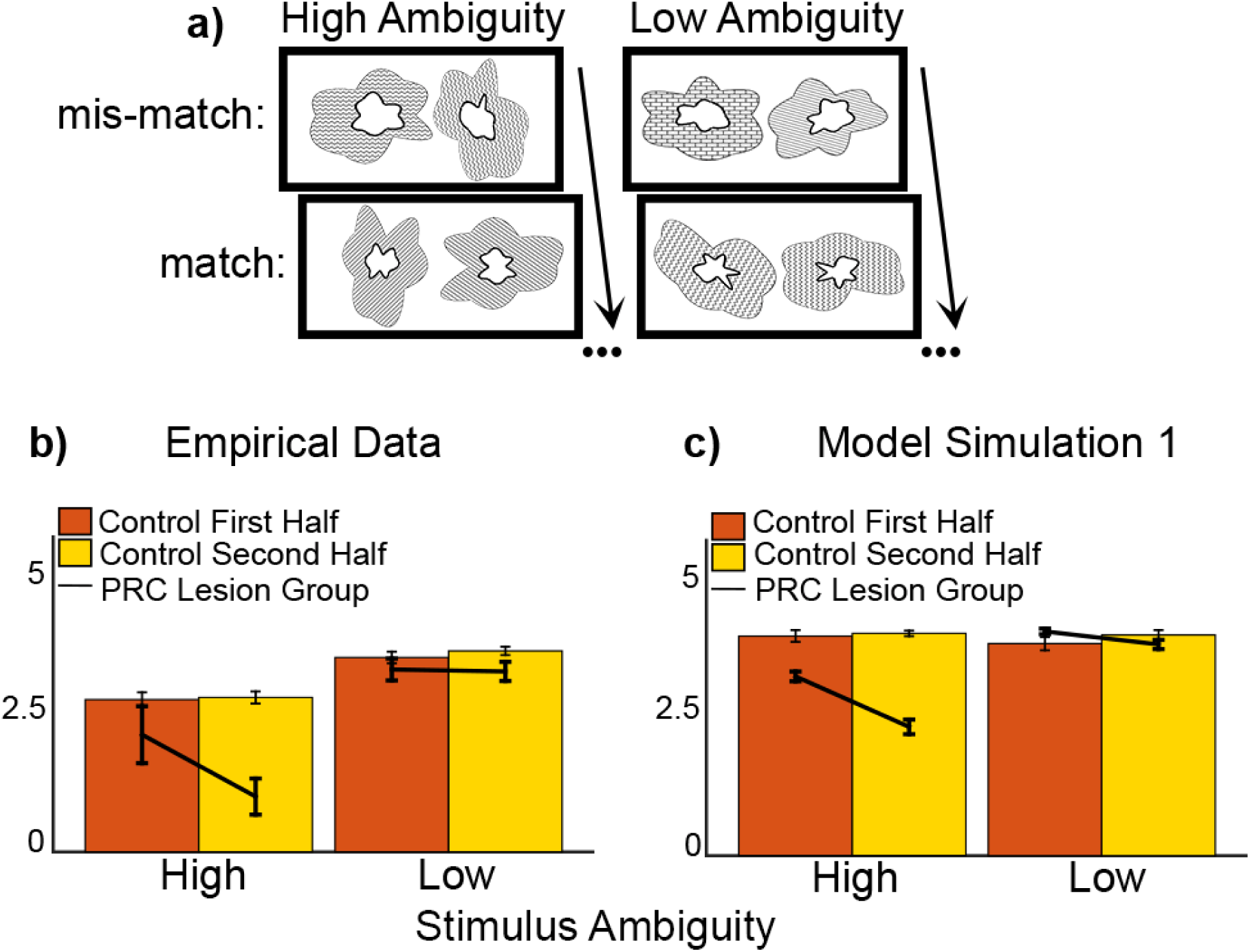
**a)** Stimuli from Experiment 3 of Barense et al. (2012). Pairs of stimuli were presented simultaneously. Each stimulus was defined by 3 features: inner shape, outer shape, and fill pattern. High Ambiguity mismatching pairs shared 2 of these features, but Low Ambiguity mismatching pairs share 0 features. **b)** Empirical data from Experiment 3 of Barense et al. (2012). Subjects with perirhinal (PRC) lesions were impaired at discriminating High Ambiguity stimuli in the second half of trials. Significance was assessed via Crawford’s t-test for each Lesion participant separately (Control n=8; Lesion n=2, Error Bars = SEM). **c)** Simulated data for Experiment 3 of Barense et al. (2012). As in the empirical data, networks with PRC lesions were impaired at discriminating High Ambiguity stimulus pairs in the second half of trials. Error Bars = SEM.

Barense et al. explained their data in terms of the representational-hierarchical account: Individuals with MTL damage lack the conjunctive representations of objects usually found in perirhinal cortex (PRC), a structure in the MTL. Objects are instead represented as a collection of features in posterior visual cortex. In the task of Barense et al., each stimulus is a unique conjunction of features, but the features comprising the stimuli repeat across trials. After viewing many items, feature-level interference renders the representations in posterior visual cortex inadequate for solving difficult (High Ambiguity) discriminations. Control subjects resolve this interference using the unique conjunction for each stimulus represented in PRC, but when PRC is damaged, performance suffers.

These results present a paradox: the *decrease* in MTL amnesics’ performance with increasing exposure to task stimuli contrasts with perceptual learning effects. Perceptual learning is often explained by assuming that experience increases the separability of stimuli, either because stimulus representations become less overlapping (Saksida, 1999; Schoups, Vogels, Qian, & Orban, 2001; Yang & Maunsell, 2004), or because the weights via which stimulus representations influence decision-making are optimized (Kumano & Uka, 2013; Liu, Dosher, & Lu, 2015). But if discrimination relies on the separability of stimulus representations, and exposure differentiates representations, then it is not clear why brain damage should reverse this effect, causing exposure to hurt performance. Put another way, even if individuals with MTL damage possess only feature representations, why should exposure should cause feature representations to become more overlapping, rather than less?

In a previous neural network instantiation of the representational-hierarchical account (Cowell et al., 2006) impairments in *recognition memory* induced by MTL damage were accounted for by considering the familiarity signal evoked by stimulus representations in the brain. Exposure to many items sharing visual features entails frequent repetition of the features. Eventually, the representations of *all* features in posterior visual cortex bear the hallmarks of familiarity, causing individuals with MTL damage (who possess only feature representations) to perceive all items as equally familiar, impairing recognition memory. Here, we invoke the same mechanism to explain the *visual discrimination* impairments reported by Barense et al. To apply this account to visual discrimination, we assume that participants use a familiarity-based comparison to decide whether two items are identical (Dai, Versfeld, & Green, 1996; Macmillan & Creelman, 2005). Although the term ‘familiarity’ provides an intuition as to how a model of memory can be applied to visual discrimination, a more accurate and theory-neutral description of the mechanism underlying *both* tasks is given by the terms ‘sharpness’ (Norman & O’Reilly, 2003) or ‘representational tunedness’ (Cowell et al., 2006).

### Resolving the Paradox: Visual Discrimination Based on Representational Tunedness

We assume that, for difficult discrimination tasks like that of Barense et al. (2012), participants search for a mismatch between the two stimuli (Dai et al., 1996). To do so, they visually scan back and forth between stimuli in a pair; if the second item appears less tuned than the item just examined, this provides evidence for a mismatch in identity. That is, the new item appears novel to the extent that it differs from the item just inspected. This forms the critical basis of the discrimination mechanism: If switching between items elicits a novelty signal (i.e., a decrease in representational tunedness) the two stimuli are judged to mismatch; if not, the two stimuli are judged to match.

Just as in the memory simulations of Cowell et al. (2006), representations in the model can saturate (i.e., reach a maximum) in terms of tunedness. In a discrimination task in which all stimuli are similar, the stimulus features appear repeatedly, resulting in all feature representations becoming highly tuned. After sufficient repetitions, the representations of all object features are equally, maximally, tuned – rendering the tunedness-based comparison inutile. In contrast, because individual objects do not repeat across trials, object-level representations (in PRC) do not saturate and so remain useful for discrimination. When conjunctive representations are compromised by MTL damage, visual discrimination is impaired. A central assumption of the representational-hierarchical framework is that memory and perception share common neural mechanisms. Our model embodies this by using a mnemonic signal – familiarity, or tunedness – to solve a visual perceptual task.

## GENERAL METHODS

### Model Architecture

We use the model of Cowell et al. (2006) with minor modifications. The network contains a PRC layer and a layer corresponding to posterior ventral visual stream (Fig. 3). Visual objects are instantiated as 8-dimensional vectors. We assume that posterior regions represent simple conjunctions of two visual dimensions, so the Posterior layer is divided into four grids: posterior grid units receive two input dimensions and combine them into a simple conjunction, termed a ‘feature’. Because PRC is assumed to represent whole objects, all eight input dimensions converge into a single 8-dimensional conjunction in the PRC layer. Thus, PRC contains unique, conjunctive representations of objects, whereas the Posterior layer represents the four 2-dimensional features separately. The Posterior layer does not input to the PRC layer; the two layers’ representations are independent. This parallel architecture greatly simplifies the model. Although the ventral visual pathway contains many serial connections (Desimone & Ungerleider, 1989), mapping the transfer of information across serial layers of the hierarchy would add extra complexity without contributing to the mechanism that accounts for the target effects. We instead opt for parsimony, choosing a simple parallel architecture that allows us to focus on our central hypothesis concerning the consequences of removing high-dimensional, conjunctive representations.

**Figure 3.**
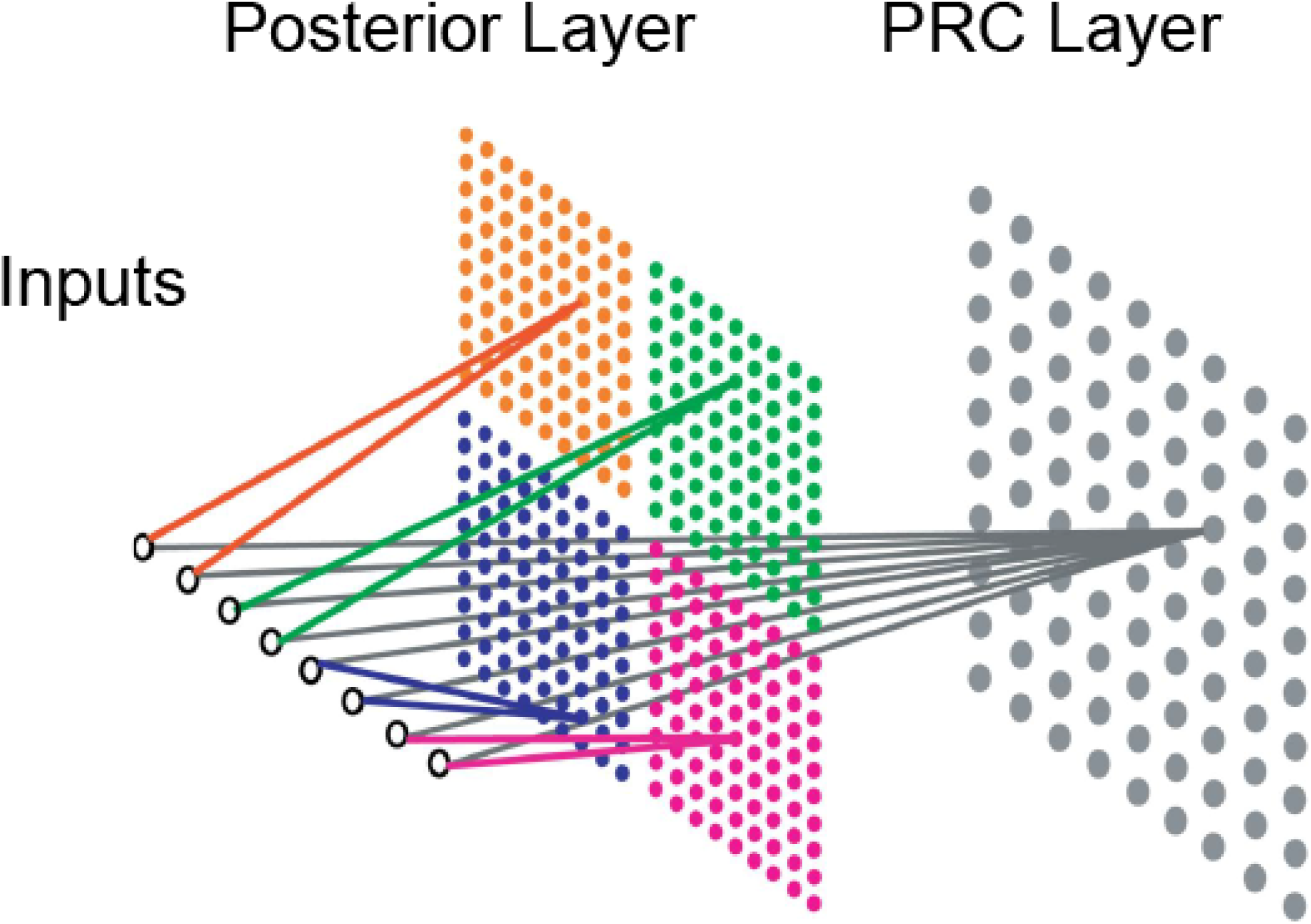
Model architecture. An object stimulus has eight input dimensions, paired into four 2-dimensional ‘features‘. The PRC layer is a single Kohonen grid, representing an object as a unique conjunction. The Posterior layer is composed of four Kohonen grids, which each represent one visual feature. Encoding and retrieval are identical on Posterior and PRC layers; the layers differ only in the complexity of representations.

All model layers are constructed from Kohonen grids, which mimic information processing mechanisms of cortex such as Hebbian learning and lateral inhibition (Kohonen, 1984). A Kohonen grid (or self-organizing map) is trained by presenting a series of stimulus inputs and incrementally adapting the weights of grid units on each presentation. During learning, stimulus representations are sharpened such that they are more tuned to a particular stimulus. Once a stimulus has been encoded, its representation on the grid is more selective: a smaller region of the grid is active, but the magnitude of that activation is greater (Fig. 4). In simulations of memory tasks, the sharpness, or ‘tunedness’, of a representation provides an index of familiarity (e.g., Cowell et al., 2006; Norman & O’Reilly, 2003).

**Figure 4.**
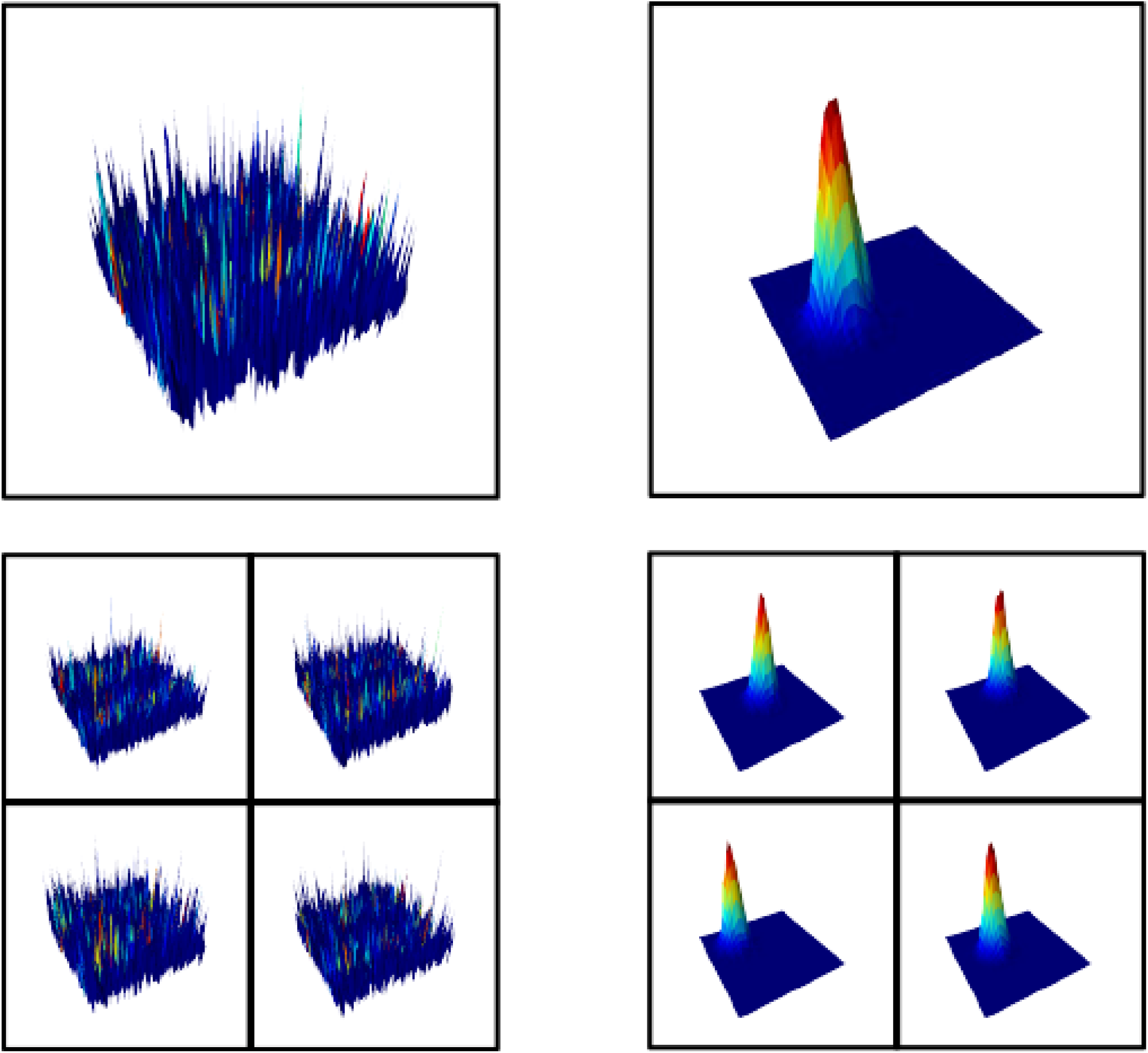
Schematic illustration of activations in the network. Left Column: Activation due to a novel stimulus (upper panel shows PRC layer, lower panel shows Posterior layer). Right Column: activation due to a stimulus that has been encoded, i.e., sampled many times. The novel stimulus elicits a broadly distributed pattern of activity across the grid, whereas the encoded stimulus elicits a highly selective activation pattern with a peak over a subset of grid units.

Each input dimension can take one of four values: 0.05, 035, 0.65, or 0.95. This scheme yields 4^8^ = 65,536 unique objects on the PRC layer, but only 4^2^ = 16 unique 2-dimensional features on each Posterior grid, reflecting a key assumption of the representational-hierarchical account: that there is a vast number of possible visual objects in the world, composed from a small number of visual elements.

### Simulating Visual Discrimination Behavior

*Fixations.* In Barense et al., participants decided whether two simultaneously presented stimuli were the same or different. Eye-tracking data revealed that, on trials ultimately declared as a ‘match’, control subjects made approximately 25 and 20 fixations at High and Low Ambiguity, respectively. These fixations were distributed across the two stimuli, as participants looked back and forth between them. Participants exhibited a higher ratio of within-stimulus to between-stimulus fixations in High Ambiguity (~1.2) than in Low Ambiguity trials (~0.6). Barense et al. conjectured that this reflected a greater tendency to bind features together in the High Ambiguity condition. Accordingly, we hypothesized that fixation ratios might contribute to task performance (e.g., sampling with a higher within:between ratio may facilitate the formation of conjunctive representations in PRC) and aligned simulation parameters with these empirical data. In the model, stimuli are ‘sampled’ via fixations. On each fixation, the network encodes the stimulus for 20 encoding cycles (see APPENDIX). During each encoding cycle, the representation of the sampled stimulus is tuned. A probabilistic rule governs how the network switches back and forth between sampling the two items in a pair, with the probability of a switch derived from the empirical within:between ratio for each experimental condition (1.2 or 0.6).

*Discrimination Decisions.* To perform a discrimination task, a network must make a ‘match/mismatch’ response; the mechanism by which it does so is as follows. On each trial, two stimuli are presented to the network, but only one stimulus is sampled (i.e., its representation ‘activated’) at a time. One stimulus is arbitrarily selected first (Item A) and the network samples it via successive fixations until switching to the other stimulus (Item B). Upon switching, the model assesses evidence for a mismatch by computing a novelty score and comparing it to a criterion. The novelty score is obtained by taking the tunedness of Item A and subtracting the tunedness of Item B. When the two items are identical, the novelty score is zero; when they are different, Item B is slightly less well tuned than Item A (because Item A has just been encoded, whereas Item B has not), yielding a positive novelty score. If the novelty score in any individual grid (any of the 4 Posterior grids or the PRC layer) exceeds the criterion, the items are declared to ‘mismatch’. If the network finds no evidence for a mismatch after this switch, fixations continue. In the next comparison, Item B serves as the previously inspected stimulus and Item A as the newly fixated stimulus. Comparisons proceed until either a mismatch is declared or the maximum number of fixations is reached, whereupon a match is declared.

We note that this mechanism disregards any information about the perceptual identity of the object and its features, which would be indexed in this model by the *location* (rather than the tunedness) of the peaks in the grid. As such, it is an unconventional mechanism for performing ‘match/mismatch’ discriminations; we return to this issue in the *Simulation 1 Discussion* and *Interim Discussion*, below.

Reaching the maximum number of fixations corresponds to the model ‘giving up’ on the search for a mismatch. Human control participants fixated longer on matching High ambiguity pairs as compared to matching Low ambiguity pairs. The study design was blocked, with High versus Low Ambiguity trials being presented in separate blocks. We assume that the blocked format allowed participants to perceive that differences between the two stimuli in a pair were subtler on High Ambiguity trials, leading participants to exercise extra caution in declaring a match, and resulting in a greater total number of fixations on matching High Ambiguity trials. We instantiate this cautiousness by setting the maximum number of fixations in the model to the values observed in control participants – 25 and 20 in the High and Low ambiguity conditions, respectively.

Finally, as in Barense et al. (2012), hit rates of 1.0 and false alarm rates of 0.0 (where ‘hit’ and ‘false alarm’ refer to declaring ‘mismatch’ and ‘match’ to a pair of mismatching stimuli, respectively) were adjusted by subtracting or adding half a trial, respectively.

*Criterion Shift.* We assume that participants adjust their decision rule as their representations adapt: if participants begin to perceive all items as more similar, they require less evidence to declare that two items are mismatching. To effect this, the criterion value of novelty required to declare a pair of items as mismatching (i.e., the *decision criterion*) is allowed to shift by equating it to the average of the novelty score on the previous 6 trials. In the present task, the assumption of a criterion shift is necessary in order to prevent an unrealistic scenario in which participants declare all pairs of items as matching, as interference accrues. However, in the absence of noise, the criterion shift would give rise to unrealistically perfect discrimination performance for all participants. Therefore, on each trial, noise drawn from the uniform distribution (±1*e* ^-6^) is added to the criterion. The properties of the noise distribution are not critical to the representational-hierarchical account; a uniform distribution was chosen for simplicity. This noise – which may encompass neural noise, attentional fluctuations, item differences, etc. – adds randomness to the decision process. In doing so, it more effectively obscures the novelty signal produced by switching between two stimuli when their tunedness difference is minimal. The starting criterion value (i.e., on Trial 1) is set to twice the maximum noise: 2*e* ^-6^.

## SIMULATION 1

We simulated the paradoxical finding that visual discrimination performance worsens with exposure to the task stimuli after damage to MTL. In *Experiment 3* of Barense et al., participants indicated whether two simultaneously presented visual stimuli were a match or a mismatch. Stimuli were trial-unique items composed of three features (Fig. 2a). In High Ambiguity mismatch trials the items shared two out of three features, whereas in Low Ambiguity mismatch trials the items shared none. Individuals with PRC damage performed like controls at Low Ambiguity, but at High Ambiguity their performance was intact initially, then fell sharply in the second half of trials (Fig. 2b).

### Simulation 1 Methods

All stimuli were trial-unique. Stimulus pairs in the Low Ambiguity condition shared no 2-dimensional features, whereas High Ambiguity pairs shared three out of four features. (The stimuli in Barense et al. contained three explicitly defined features, but we used four features for consistency with Cowell et al. (2006); the total number of features does not qualitatively affect simulation results). Because the representational-hierarchical account assumes that objects are composed from a limited pool of visual features, there are many possible unique objects, but the features comprising them appear repeatedly. For this simulation, we further constrained the feature set to reflect the high feature-overlap among the stimuli of Barense et al. by constructing four-featured objects using only six of the sixteen possible 2-dimensional features for each Posterior grid. This yielded 6^4^ = 1296 unique objects.

Networks were initialized and pre-trained (see APPENDIX) before performing visual discrimination in two conditions: High and Low Ambiguity. As in Barense et al., each condition contained 36 ‘match’ and 36 ‘mismatch’ trials. Control networks comprised both posterior and PRC layers, whereas ‘PRC Lesion’ networks possessed only a Posterior layer. We simulated 48 networks in each group, corresponding to 6 networks per human control participant in Barense et al. (2012).

### Simulation 1 Results and Discussion

Networks with no PRC layer, like humans with PRC damage, were impaired relative to controls at High but not Low Ambiguity (Fig. 2c), and the impairment was worse in the second half of trials. We do not report statistics on simulated data because significance scales arbitrarily with the number of networks run. Instead, we focus on qualitative patterns, which match those of the patient data, including the critical interaction between Lesion Group, Ambiguity and First/Second Half.

The simulation of a lesion-induced deficit in discriminating High Ambiguity stimuli in the second half of trials hinges on three assumptions: (1) participants solve the task using a tunedness-based differencing rule; (2) the stimuli contain many low-level features that repeat over trials so that eventually all stimulus features are highly tuned; (3) the stimuli are represented in PRC as whole conjunctions but in posterior regions as individual features. Together, these assumptions provide that, following PRC damage, discrimination performance is impaired once all features are maximally tuned.

Consider a Lesioned network, in which performance relies on feature representations because it possesses only a Posterior layer. At the start of the task, individual features are not highly tuned. On each new trial, as the network encodes the first stimulus, the features of that stimulus are tuned. When the network switches to the second stimulus, if it is not identical to the first its features appear novel and a mismatch is correctly declared. However, after many trials, all features have been repeatedly encoded by the network. Now, at the start of a new trial, there can be no increase in tunedness of the feature representations when the network inspects the first stimulus. Next, when the network switches to the second stimulus, even if that stimulus differs in identity from the first, its features are just as tuned as those of the first, and the tunedness-based differencing rule cannot reliably detect a mismatch.

The feature-level interference has more effect at High than Low Ambiguity for two reasons. First, networks perform more encoding on any given mismatch trial in the High than in the Low Ambiguity condition. This is because the network spends longer searching for a difference between the two stimuli on High Ambiguity trials than on Low Ambiguity trials. The longer search time is produced by the higher ratio of within:between fixations on High Ambiguity trials (see Fixations.). In searching longer for a mismatch, more encoding occurs, and stimulus representations undergo more tuning. Consequently, the tunedness of features rises more steeply across trials in the High Ambiguity condition. Although – as Barense et al. claimed – the use of a higher within:between fixation ratio may be useful for binding features into a conjunction, that strategy ultimately proves detrimental to the lesioned model: inspecting each stimulus more closely leads to faster build-up of interference.

Evidence that humans with brain damage nevertheless adopt this disadvantageous strategy is provided by Erez, Lee, and Barense (2013), in which patients with PRC damage exhibited the same viewing patterns as control subjects. The second reason that lesioned networks’ performance deteriorates faster at High Ambiguity is that mismatching High Ambiguity pairs share three out of four features, whereas Low Ambiguity pairs share no features. In seeking a mismatch, the network searches for any pair of features across the two stimuli that differ. In Low Ambiguity pairs, there are four mismatching features therefore a network has four opportunities to discover a feature that has not yet reached maximum tunedness, which can be used to declare the two items a mismatch. In High Ambiguity pairs, there is only one mismatching feature and so the chance of discovering a mismatch is greatly reduced.

Performance in control networks, by contrast, is maintained throughout the task, because they possess conjunctive representations in the PRC layer. Individual stimuli are trial-unique (i.e., whole conjunctions are never repeated) so whole-object representations in PRC never reach maximum tunedness. At the start of each new trial, the PRC representation for the first stimulus inspected always becomes more selective as it is inspected. When the network switches to the second stimulus, if the second differs from the first, the second will be less tuned and the pair will be declared to mismatch.

This mechanism for performing visual discrimination is radically different from traditional conceptions of how same/different (‘match/mismatch’) judgments are made. Traditional theories of discrimination typically assume that match/mismatch responses are based on the perceived identity of the objects or their features – when the identity is the same, a match response is declared. The alternative, ‘tunedness’-based mechanism of the present model is perhaps counter-intuitive: for example, does it lead to the implausible prediction that two completely physically different stimuli will be called ‘same’ if they have the same tunedness? In practice, the answer is ‘no’. If two items differ in identity, the chance that their familiarity values – which are determined by the network’s prior experience with these and other items – will be similar enough to elicit a ‘match’ response is vanishingly small (except in the posterior layer after sufficient interference). Indeed, this phenomenon is the crux of the mechanism: the model exploits the fact that different objects elicit different levels of familiarity, interpreting the differing levels of familiarity as indicative of different objects. This phenomenon breaks down in exactly the scenario that causes impairments in human subjects: when only posterior representations are available and interference has built up. Moreover, as explained in the *Interim Discussion* below, we do not suggest that the tunedness-based discrimination heuristic is used by human participants for *all* discrimination tasks (including those in which paired items differ greatly in identity), rather only for very difficult discriminations employing highly similar stimuli.

These simulations demonstrate an explicit mechanism by which perceptual interference can cause MTL amnesics to show worsening visual discrimination with increasing stimulus exposure. However, in Barense et al.’s *Experiment 3*, perceptual interference was incidental rather than explicitly manipulated – its effects were examined by comparing the first and second halves of the study, which confounded degree of interference with order of presentation. Thus, although interference was hypothesized to account for the patients’ greater impairment in the second half of trials, a potential alternative explanation was that patients grew relatively more fatigued than controls as the task wore on, rendering their performance in the harder (High Ambiguity) condition more impaired. To test directly the hypothesis that perceptual interference is central to MTL patients’ deficits, Barense et al. conducted another study – *Experiment 4* – in which perceptual interference was explicitly manipulated.

## SIMULATION 2

In *Experiment 4*, subjects completed three blocks in strict order: Low Interference 1, High Interference, and Low Interference 2. The High Interference block contained 88 trials (44 match, 44 mismatch) identical to the High Ambiguity trials of *Experiment 3*, in which pairs of abstract stimuli shared two out of three features. In Low Interference blocks, High Ambiguity and photographic trials were interleaved: each of 30 High Ambiguity trials (15 match, 15 mismatch) was followed by two trials containing a pair of color photographs of real-world objects (58 trials in total; 29 match, 29 mismatch). Critically, performance was assessed only on every third trial, which was always a High Ambiguity trial. Photographic stimuli shared few low-level features with the abstract stimuli of High Ambiguity trials (Fig. 5a). Therefore, Low Interference blocks entailed less feature-level interference than High Interference blocks, and MTL patients were predicted to be less impaired at Low Interference. *Experiment 4* replicated the results of *Experiment 3*: MTL-damaged patients showed impaired discrimination at High but not Low Interference, even in the second Low Interference block (Fig. 5b). This suggested that the impairment seen in *Experiment 3* was caused by the cumulative effect of perceptual interference within a block, rather than increasing fatigue in the MTL patients.

**Figure 5.**
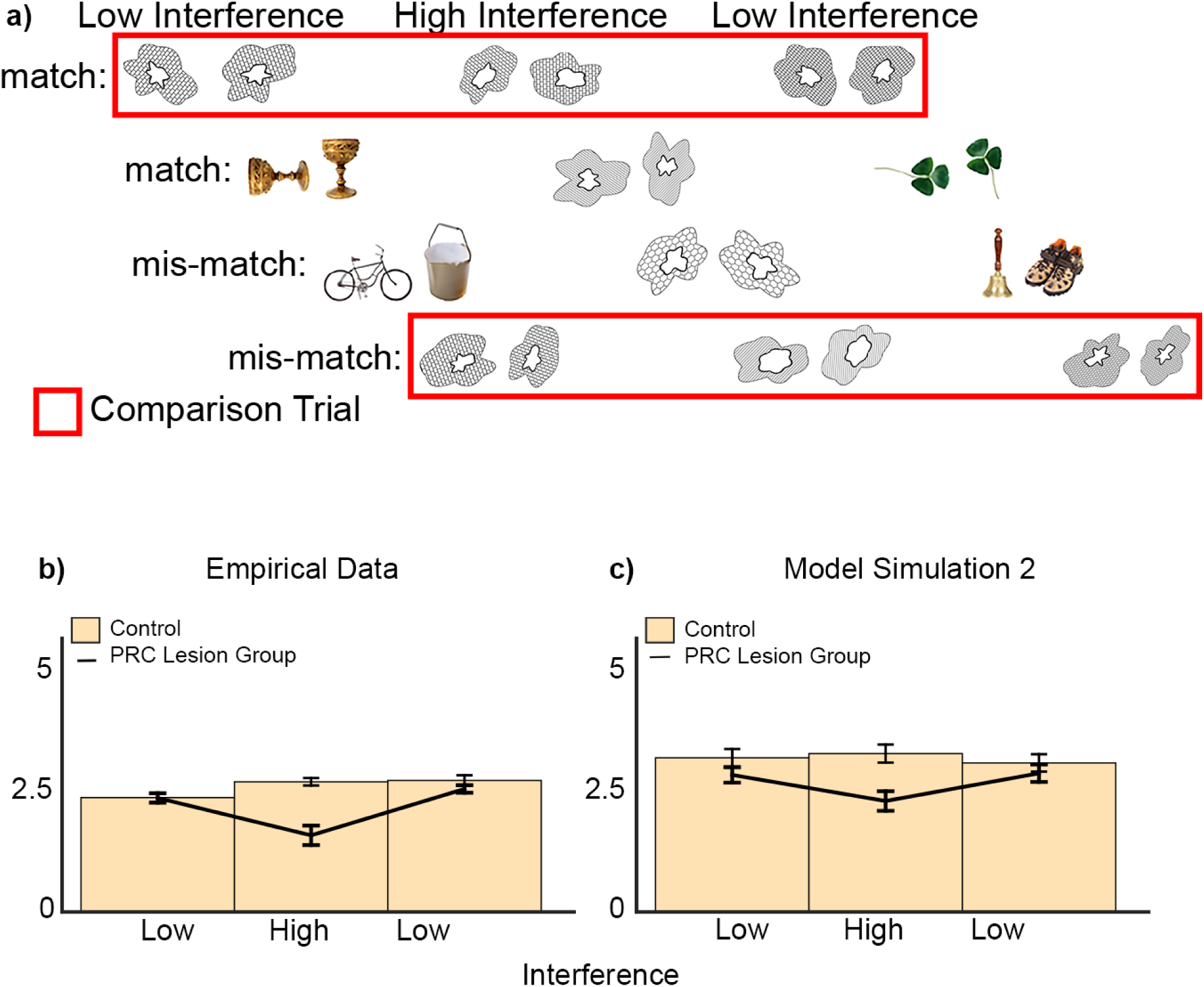
**a)** Stimuli from Experiment 4 of Barense et al. (2012). Low Interference blocks used photo stimuli in 2/3 of trials, which shared few features with the abstract stimuli used on critical comparison trials. High Interference blocks used abstract stimuli on every trial. **b)** Empirical data from Experiment 4 of Barense et al. (2012). **c)** Model simulations for Experiment 4. Error Bars = SEM.

### Simulation 2 Methods

As in Simulation 1, we modeled abstract stimuli by using six of the sixteen possible ‘features’ on each Posterior grid, to construct four-featured stimulus wholes with high feature-overlap. Reflecting the assumption of Barense et al. that abstract stimuli shared few low-level features with photographic stimuli, we used the remaining ten features (i.e., an independent set) to construct photographic inputs.

Networks performed three discrimination blocks, each containing 88 trials. Every third trial in all blocks was a critical comparison trial, in which the two stimuli were abstract stimuli (15 matching, 15 mismatching). On mismatching critical comparison trials, the stimuli shared two out of three features. In High Interference blocks the remaining 58 trials contained extra pairs of High Ambiguity abstract stimuli. In Low Interference blocks the remaining 58 trials contained pairs of photographic stimuli. As in Barense et al., in both High and Low Interference blocks, performance was judged only on critical comparison trials: the trials occurring at every third position. The key difference between High and Low Interference was that, for Low Interference, the trials interposed between critical trials contained items sharing no features with critical-trial stimuli whereas, for High Interference, interposed trials contained items similar to critical-trial stimuli.

### Simulation 2 Results and Discussion

Mirroring the data from MTL amnesics, lesioned networks were unimpaired relative to controls in the Low Interference blocks, but impaired in the High Interference block (Fig. 5c).

The same mechanism that impaired lesioned networks in Simulation 1 drives the impairments in Simulation 2. In the High Interference condition, because all trials contain the same class of stimuli, stimulus features appear repeatedly and posterior feature representations reach maximum tunedness. Once this saturation occurs, the tunedness of posterior, feature-based representations cannot increase substantially when the network inspects a new stimulus at the start of a trial. Consequently, a network with no PRC layer cannot detect novelty (a drop in tunedness) upon switching to the other stimulus in the pair, and a tunedness comparison no longer discriminates the two stimuli. In contrast, in the Low Interference condition, two-thirds of trials contain photographs composed of different features than the critical-trial stimuli. The critical-trial stimulus features repeat too infrequently for the tunedness of their representations to saturate, and lesioned networks remain unimpaired.

Notably, the discrimination scores of lesioned networks improved in the final Low Interference block, relative to the previous High Interference block. This improvement suggests a release from built-up interference, despite the fact that – to mirror the cumulative effects of fatigue or interference experienced by MTL patients – networks’ weights were *not* reset to their initial values between blocks. The improvement can be attributed to the interposition of stimuli composed of very different visual features, on two-thirds of trials. That is, because the posterior layer was required to alternate between encoding the features of the abstract stimuli (on critical trials) and a very distinct set of features of photographic stimuli (on interposed trials), the representations of abstract stimulus features became partially ‘detuned’ between critical trials, such that the tunedness-based differencing rule for discriminating two abstract stimuli was effective, even based only upon feature representations.

## INTERIM SUMMARY: ACCOUNTING FOR VISUAL DISCRIMINATION

Barense et al. (2012) reported a striking perceptual deficit in patients with MTL damage: the accumulation of perceptual experience impairs visual discrimination. This result is paradoxical because perceptual discrimination typically improves with exposure to the stimuli. Barense et al. argued that MTL-lesioned patients suffer from accumulated feature-level interference, which − in the absence of conjunctive MTL representations − cannot be overcome by feature representations in posterior visual cortex. Although we concur with this explanation, we suggest that it is incomplete.

Standard theories of perceptual learning claim that experience improves discrimination performance by reducing the overlap between stimulus representations (i.e., training increases representational separation). In such theories, the assumed mechanism for visual discrimination is that discriminability is proportional to the overlap between representations (Saksida, 1999; Schoups et al., 2001). But this mechanism cannot account for the data of Barense et al.: if exposure separates representations, even feature-based discrimination should improve with exposure, because even feature representations should become less overlapping with exposure. To explain why the performance of MTL patients in Barense et al. worsened after exposure, a theory based on representational overlap would require the counter-intuitive assumption that – although cortical representations of stimuli underlying perceptual learning typically become less overlapping with exposure (e.g., Jenkins, Merzenich, Ochs, Allard, & Guíc-Robles, 1990) – posterior feature representations become more so.

In the account provided by Simulations 1 and 2, we eschew representational overlap as the mechanism for visual discrimination. Instead, we take the explanation offered by Barense et al. – that amnesics suffer from compromised conjunctive MTL representations – and combine it with a less commonly invoked discrimination mechanism: a familiarity-based differencing rule, which capitalizes on differences in the tunedness of representations caused by moment-to-moment encoding. Under this account – as in prior instances of the representational-hierarchical account applied to memory (Cowell et al., 2006) – representations of features, but not conjunctions, reach maximum tunedness as interference accrues. Thus, after MTL damage, perceptual experience impairs visual discrimination.

Two aspects of the mechanism for visual discrimination require clarification. First, although we simulate just two layers of the ventral pathway, this comprises only a subset of the full hierarchy of representations in the brain, which includes lower-dimensional layers prior to the model’s Posterior layer and higher-dimensional layers, such as hippocampus, after PRC. Other tasks with different representational requirements may require other layers (Cowell, Bussey, & Saksida, 2010). For example, a discrimination task involving whole objects that repeat would require hippocampal representations capable of combining objects with contextual or temporal information, to shield participants from object-level interference that cannot be resolved in PRC (see *Simulation 5*). Second, we do not suggest that a tunedness-based differencing rule is used in all discrimination tasks. Dai et al. (1996) point out that a differencing rule is optimal (and adopted) when familiarity signals for the to-be-discriminated stimuli are highly correlated. Familiarity signals would be less correlated in easy discrimination tasks in which the stimuli differ in terms of salient features like color. Such tasks could be solved by a more traditional discrimination mechanism that assesses items’ perceptual identity, rather than their familiarity. In our model, perceptual identity corresponds to the location of a representation in the grid, rather than its tunedness, and a traditional discrimination mechanism would involve assessing the representational overlap between two stimuli. The alternative, tunedness-based differing rule that is critical to producing the pattern of data observed in the MTL patients of Barense et al. (2012) is thus an unconventional mechanism, and it likely only applies when differences in perceptual identity between stimuli are small and familiarity signals are highly correlated. In other words, we do not doubt that conventional, perceptual-identity-based discrimination mechanisms are employed by human participants in many discrimination tasks; we suggest only that participants use a tunedness-based differencing rule for discriminating highly similar stimuli in very difficult tasks.

In sum, the model of Cowell et al. (2006) – originally developed to explain recognition memory – accounted for deficits in perceptual discrimination observed in patients with PRC damage. Critically, the mechanism by which lesioned networks are impaired at visual discrimination is the same mechanism that causes deficits in recognition memory. In both cases, lesioning the PRC layer removes the conjunctive representations that ordinarily shield the network from feature-level perceptual interference. In both cases, the remaining posterior, feature-based representations reach an asymptotic level of tunedness, rendering them incapable of solving the task.

## A UNIFIED MODEL OF MNEMONIC AND PERCEPTUAL DEFICITS

Barring some minor modifications, the network that accounted for visual discrimination in Simulations 1 and 2 retained the architecture and parameters of the recognition memory model of Cowell et al. (2006). Nonetheless, to verify that the modifications did not qualitatively change the predictions for recognition memory, we resimulated the three key findings concerning deficits in recognition memory caused by PRC lesions: (1) the deficit is delay-dependent: it worsens as the retention interval increases; (2) the deficit is exacerbated by increasing the length of the list of studied items; and (3) recognition memory for repeatedly-presented stimuli is not impaired by PRC lesions.

### General Methods for Simulating Recognition Memory

Following Cowell et al. (2006), these deficits were accounted for by simulating an object recognition task akin to the spontaneous object recognition (SOR) and delayed non-match-to-sample tasks (DNMS) used in animals (Ennaceur & Delacour, 1988; Mishkin & Delacour, 1975). In these tasks, subjects are presented with a list of items (the ‘study’ or ‘sample’ phase). After a retention delay, a copy of each studied item is presented, paired with a novel item. In the SOR task, healthy animals spontaneously spend more time exploring the novel than the familiar item, yielding a ‘recognition score’. In DNMS, the animal is rewarded for choosing the item that was not previously encountered. Both tasks require the detection of a difference in familiarity between the novel and familiar items.

To simulate recognition memory we follow the protocol of Cowell et al. (2006). Accordingly, aspects of Simulations 1 and 2 that were intrinsic to visual discrimination behavior (e.g., fixations, criterion shifts) are not included. Briefly, a pre-trained network encodes a list of stimuli during the study phase and judges the familiarity of studied and novel objects at test. In tasks involving a delay between study and test, we assume that forgetting is caused by visual interference, which is simulated by presenting a series of randomly selected stimuli. In the test phase, the network is presented with the list of sample stimuli, each paired with a novel item. An index of familiarity –‘Tunedness’ – is calculated for both sample and novel items, and a recognition score is derived by combining them into a normalized difference score (Appendix, Equation 5). A higher recognition score indicates greater familiarity of the sample than the novel item, i.e., better recognition memory performance. Empirical SOR and DNMS tasks typically use a diverse sample of everyday objects as stimuli and so we constructed stimuli using all possible visual features (16 per posterior grid; 16*4 = 64 in total) to reflect the greater variation among stimuli than in Barense et al. (2012). As in Simulations 1 and 2, we assess model performance by examining qualitative trends in the simulation results.

## SIMULATION 3: DELAY-DEPENDENT DEFICITS IN RECOGNITION MEMORY

In a recognition memory task, increasing the delay between study and test is assumed to increase the load on memory, and so manipulating the delay length varies the extent to which a putative memory system is taxed (Cowell et al., 2006; Gaffan, 1974). Animals with PRC lesions exhibit worsening recognition memory deficits with increasing delay, strongly implicating PRC in recognition memory (Buffalo, Ramus, Squire, & Zola, 2000; Buffalo, Reber, & Squire, 1998; Eacott, Gaffan, & Murray, 1994; Malkova, Bachevalier, Mortimer, & Saunders, 2001; Meunier, Bachevalier, Mishkin, & Murray, 1993; Mumby & Pinel, 1994).

### Simulation 3 Methods

There were four trials, each comprising three phases: study, delay and test. Delay was simulated by presenting interfering stimuli, which were randomly selected, with replacement, from the pool of all possible stimuli (65,536 objects), and presented to the network for one encoding cycle each. More interfering stimuli corresponded to a longer delay; delays ranged from 0 to 8000 stimuli.

### Simulation 3 Results and Discussion

Removing the PRC layer of the network caused deficits in recognition memory that worsened with lengthening delay (Fig. 6a). The mechanism underlying the deficit is shared with Simulations 1 and 2 of the Barense et al. data. The interfering items encountered during a delay are composed of a limited number of features: individual features appear repeatedly, whereas the unique conjunctions of features corresponding to whole objects do not. Consequently, feature-based representations on the Posterior layer are rendered familiar by interference, whereas conjunctive PRC representations are not. In the test phase, after a delay, the Posterior representations of both the sample and the novel object appear familiar, because all features have appeared repeatedly. Lesioned networks, which possess only the Posterior layer, can no longer adequately discriminate the items on the basis of tunedness.

**Figure 6.**
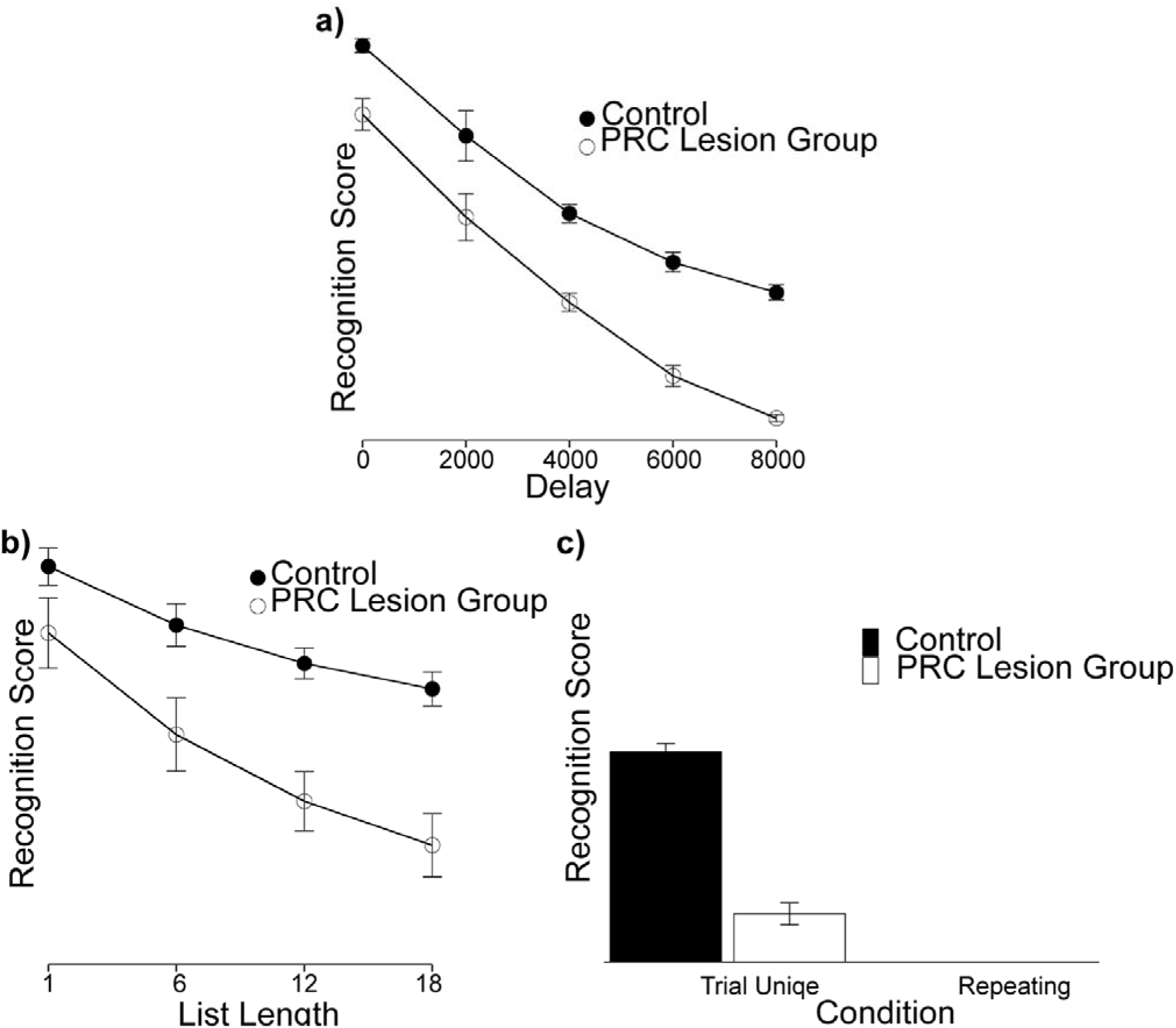
**a)** Simulation of a delay-dependent deficit in recognition memory (Simulation 3). The impairment in recognition memory is evident in the lower recognition scores for Lesioned networks, and this impairment increases as the delay between study and test increases. Abscissa indicates number of interfering stimuli sampled during the delay. **b)** Simulation of the effects of list length on recognition memory (Simulation 4). Recognition scores decrease as the sample list length increases, and the rate of decrease is faster rate for networks without a PRC layer. **c)** Simulation of the effects of trial-unique versus repeated stimuli (Simulation 5). Recognition scores are lower in lesioned networks when sample and novel stimuli are trial-unique. There is no group difference in recognition memory when stimuli are repeated because scores in both groups are equally poor. These data replicate Cowell et al. (2006). Error Bars = 95% CIs.

## SIMULATION 4: THE EFFECT OF LIST-LENGTH

Increasing list length, like increasing delay, is assumed to increase the memory load in a recognition memory task (Gaffan, 1974). Therefore a second piece of evidence that PRC is critical to recognition memory is that longer sample lists exacerbate the deficit caused by PRC lesions (Eacott et al., 1994; Malkova et al., 2001; Meunier et al., 1993).

### Simulation 4 Methods

We used lists of 1, 6, 12, or 18 pairs of unique stimuli. The sample and novel items in each pair shared no features, but features appeared repeatedly across items within a list. As in animal studies of the list-length effect, networks encoded all sample items in the list before proceeding to the test phase. To simulate simultaneous retention of all sample items in memory, network weights were not reset between encoding each successive sample item in the list.

### Simulation 4 Results and Discussion

The recognition memory impairment in lesioned networks increased as a function of list length (Fig. 6b). As in Simulation 3, lesioned networks are impaired by the accumulation of feature-level interference: with longer lists, networks are more likely to repeatedly encounter all possible visual features during the study phase; at subsequent test, feature-based representations of all objects – including novel items – appear familiar, rendering the sample and novel items indiscriminable.

## SIMULATION 5: TRIAL-UNIQUE VERSUS REPEATED STIMULI

PRC lesions impair recognition memory only when stimuli are trial-unique; when stimuli are presented repeatedly, animals with PRC damage are unimpaired (Eacott et al., 1994). In the extreme version of repeated-items recognition, the stimulus set comprises only two stimuli, which appear on every trial. Under these conditions, PRC-lesioned animals are unimpaired and, in addition, neither lesioned nor control animals perform well at a short study-test delay of 30 seconds.

### Simulation 5 Methods

Two sets of 30 pairs of stimuli were created: in the ‘Trial-unique’ set, no item appeared more than once; in the ‘Repeating’ set, the same pair of items appeared 30 times, with sample and novel status assigned randomly on each trial. In both conditions, networks completed 30 trials, with each trial comprising sample presentation, a brief delay (200 interfering items, 1 encoding cycle each) and test phase. To allow the cumulative effects of stimulus type (trial-unique versus repeated) to influence performance, network weights were not reset between successive trials.

### Simulation 5 Results and Discussion

Lesioned networks were impaired when stimuli were trial-unique, but when stimulus items were repeated the recognition scores of lesioned and control networks did not differ (Fig. 6c). The poor performance even in control networks for repeated-items recognition illustrates an important tenet of the representational-hierarchical account of cognition: that different tasks require conjunctive representations at different levels of the hierarchy. In the delay-dependent and list length simulations above, repeatedly occurring features created feature-level interference that was resolved by conjunctions in PRC. In repeated-items recognition, whole objects occur repeatedly: now, even the PRC layer suffers from interference, in that all objects possess highly tuned representations. The resolution of interference now requires an even higher-dimensional layer (not simulated here), in which objects are combined with time or context to form more complex conjunctions that enable recency judgments. As argued in Cowell et al., 2006, a candidate brain region is the hippocampus.

## GENERAL DISCUSSION

The goal of this study was to investigate whether deficits in both memory and perception caused by MTL damage can be accounted for by the same model employing a single mechanism.

Using the model of recognition memory from Cowell et al. (2006), we simulated the impaired performance of patients with PRC damage on two visual discrimination tasks (Barense et al., 2012). We then replicated the recognition memory simulations of Cowell et al. (2006) using model parameters identical to those of the discrimination simulations. To our knowledge, this represents the first computational model to explicitly simulate both mnemonic *and* perceptual deficits caused by PRC damage using a unified architecture and a common mechanism.

There exist numerous models of the role of MTL structures in memory (e.g., Bogacz, Brown, & Giraud-Carrier, 2001; Cowell et al., 2006; Linster & Hasselmo, 1997; Marr, 1971; McClelland, McNaughton, & O’Reilly, 1995; Norman & O’Reilly, 2003; Treves & Rolls, 1994). More recently, theoretical accounts of how MTL structures contribute to perception have been proposed (Bussey & Saksida, 2002; Clark & Maguire, 2016; Cowell et al., 2010; Elfman, Aly, & Yonelinas, 2014; Yonelinas, 2013). A detailed comparison of the present model to other formal models of PRC function can be found in Cowell (2012). Here we discuss one model that, to our knowledge, is the only other formal account of deficits in both perception and memory following MTL damage (Elfman et al., 2014). Elfman et al. propose that the hippocampus contributes to perception in a graded manner whenever high-resolution representations of relational information are required, and to memory in a thresholded manner whenever retrieval based upon a partial cue necessitates pattern-completion. The model we present contrasts with Elfman et al. (2014) in two key ways. First, Elfman et al. address the effects of focal hippocampal damage, whereas we address the effects of focal perirhinal damage.

Second, Elfman et al. postulate distinct processes in perception versus memory, localized to distinct hippocampal subregions: CA1 is the primary locus of the graded perceptual signal, whereas CA3 is uniquely suited to providing the ‘attractor’ representations required for pattern completion. In contrast, the present model deliberately avoids mapping processes onto neuroanatomical regions, instead explaining mnemonic and perceptual tasks with a single mechanism that operates on common representations.

By doing so, the model not only provides a parsimonious account of PRC function, but also raises a deeper question about cognition in the brain. The model claims that more than one cognitive function – in this case, recognition memory and visual discrimination – can depend on the same brain structure. When this occurs, should that structure’s function be characterized as dichotomous, as in Elfman et al., or should we reconsider how its function is defined? We argue for a reconsideration.

Consider the present study: we simulated two tasks, one traditionally defined as a perceptual task (the discrimination of simultaneously presented stimuli, without the need to retain information while stimuli are absent), and the other as a memory task (requiring the retention of information over a delay). One might interpret the simulation results as suggesting that the PRC can support two distinct cognitive functions. On the other hand, the mechanism that accounted for both tasks involved a judgment of familiarity, or novelty detection process, which is traditionally associated with memory (Bogacz et al., 2001; Mandler, 1980). So one might instead interpret the model as suggesting that the visual discrimination task of Barense et al. was in fact a memory task, and that perirhinal cortex specializes in making familiarity judgments. We reject both interpretations, advocating a very different alternative: that cognition in the brain is best explained in terms of representations and representational changes. Accordingly, traditionally intuitive labels for cognitive processes – such as *memory* and *perception*, or *familiarity* and *recollection* – should be eliminated from accounts of cognitive function. Although the term ‘familiarity’ describes intuitively how representations in the model change with experience, a more accurate description is given by ‘representational tunedness’. Exposure to a stimulus causes its representation to become tuned; the relative tunedness of two representations provides a means of discriminating the items. Just as we could describe this as ‘novelty detection’ in the visual discrimination task (proposing a mnemonic mechanism for a perceptual task), we could call it ‘perceptual learning’ in the recognition memory task (a perceptual mechanism for a memory task). The most accurate characterization of the model avoids process-based labels altogether, replacing them with a parsimonious, single-system account in terms of representations.

In sum, we present a unified account of the mnemonic and perceptual deficits caused by PRC damage, in which the contribution of a brain region to cognition is determined by the representations that it contains. Cognitive functions are realized through operations upon those representations, and are influenced by changes to those representations. Representational changes can be critical to the performance of a cognitive task, or they can be disadvantageous. In the visual discrimination and recognition memory tasks simulated here, representational changes in PRC critically support performance in healthy participants, but representational changes in posterior visual cortex disrupt performance when a person has PRC damage. Although a complete understanding of MTL function – to include decision-making, imagination, and spatial navigation – requires much future investigation, we suggest that a representational approach to theory-building may prove fruitful in this endeavor.

## ACKNOWLEDGEMENTS

We thank Morgan Barense and Lok-kin Yeung for sharing their data, and Morgan Barense for comments on a draft of the manuscript. This work was supported by NSF grant 1554871 to R.A.C.

## APPENDIX

**Initialization and Pre-training.** Each grid (four in the Posterior layer and one in the PRC layer) contains 200×200 nodes whose weights are initialized to random values from 0-1. For all simulations, networks are pretrained for 500 cycles according to the learning rule,

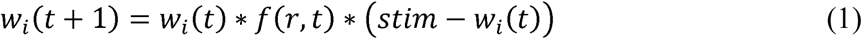

in which,

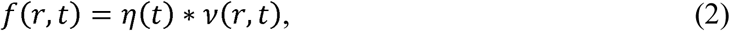

where *w*_*i*_ refers to the weights of node *i*, *t* is the current cycle, *stim* is stimulus input, *η(t)* is the learning rate, *r* is the city-block distance of node *i* from the most strongly active (winning) node, and *ν(r,t)* is a neighborhood function that scales the learning rate. Equation 1 ensures that learning on a given cycle brings a node’s weights closer to the stimulus input being presented. Equation 2 ensures that the degree to which a node’s weights are moved closer to the stimulus decreases with distance from the winning node. In the pre-training phase, both *η* and *ν* decrease with each cycle. The neighborhood function *ν(r,t)* is defined by a Gaussian function:

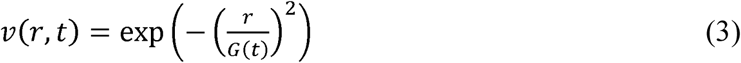

where, *G(t) = 0.5 + 10t^-B^*, and *B* is a constant determining the rate of shrinkage of the neighborhood function. The learning rate decreases as *η(t) = t ^-A^*, where the constant *A* determines the rate of decrease. On each pre-training cycle, the network is exposed to a different, unique stimulus, and its weights are updated. Because the weights of a node determine the stimulus to which it is best tuned, pre-training a Kohonen grid with a shrinking neighborhood function globally organizes the nodes, such that nodes near each other are tuned to similar stimuli (Kohonen, 1984). In our model, pre-training reflects the general, pre-experimental visual experience of participants. Because all grids in the network operate independently during experimental simulations, grids were pre-trained independently.

In all simulations, networks begin each task condition (e.g., Low Ambiguity versus High Ambiguity in Simulation 1, or 0 delay versus 2000 delay in Simulation 3) by assuming the weight state reached at the end of pre-training, unless otherwise indicated.

**Encoding a Stimulus.** In Simulations 1 and 2, stimuli are encoded during each trial as the network fixates on one stimulus at a time. Each fixation of a stimulus entails encoding it for 20 cycles, following equations 1, 2 and 3, except that *η* and *ν* are constants fixed at the values of the final pre-training cycle (*η = η(500); G = G(500)*). After a between-stimulus fixation, the network calculates a novelty score, which is the tunedness of current stimulus subtracted from the tunedness of the other stimulus. If the novelty score exceeds criterion (see main text), the stimuli are declared to mismatch and a new trial begins. If a mismatch is not declared, encoding proceeds on the current stimulus and fixations continue until either a mismatch is declared or the maximum number of fixations is exceeded.

In Simulations 3, 4 and 5, a sample stimulus is encoded following Equations 1, 2 and 3 (with fixed *η* and *ν*) for 500 encoding cycles. This reflects the encoding that would be achieved through multiple fixations, but fixations are not critical to the mechanism and so are not modeled. Because trials in Simulations 1 and 2 include multiple fixations with 20 cycles per fixation, all simulations use a similar number of encoding cycles per trial. Qualitative patterns in the results do not depend on the exact number of encoding cycles.

**Measuring Tunedness.** In Simulations 1 and 2, the ‘representational tunedness’ (intuitively, ‘familiarity’) of both stimuli is assessed whenever the network switches from fixating one stimulus to the other. In Simulations 3, 4 and 5, the tunedness of both stimuli is assessed during the test (choice) phase. We define tunedness according to the pattern of activation elicited by a stimulus. Activation *a* of node *i* is determined by the sigmoid function:

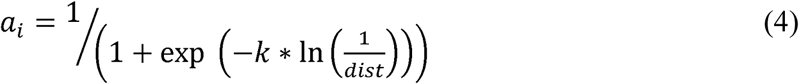

where *k* is a constant determining the steepness of the sigmoid function, and *dist* is the mean squared error between a node’s weights *w*_*i*_ and the stimulus input vector.

The tunedness, *T*, of a stimulus is calculated separately in each grid. Tunedness is the activation of the peak (the summed activation of the winning node and its nearest 4 neighbors) divided by the summed total activation of the grid. Tunedness is measured separately in each grid in the network, yielding a single, object-level tunedness score from the PRC layer and four separate feature-level tunedness scores in the Posterior layer. When comparing two stimuli in Simulations 1 and 2, posterior tunedness scores are compared separately for each pair of features. Stimulus representations are not updated during the choice phase of recognition memory simulations (Simulations 3-5).

**Recognition Score.** For Simulations 3, 4 and 5, the recognition score, *R*, is calculated as:

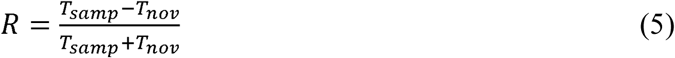

where *T*_*samp*_ is the tunedness of the sample stimulus representation, and *T*_*nov*_ is the tunedness of the novel stimulus. *R* is calculated using *T*_*samp*_ and *T*_*nov*_ values that are averaged over the separate grids of the network. In Control networks, which possess a PRC grid, *T* is computed by first averaging across all four posterior *T* values, then taking the mean of the Posterior and PRC *T* values. In Lesioned networks, *T* is the average of the Posterior *T* values.

**Parameters.** In all simulations, *k = 0.08*, *B = 0.3*, and *A = 0.6*. As in Cowell et al. (2006), we simulated 6 networks per group in the recognition memory simulations (Simulations 3, 4 and 5). In the simulations of Barense et al., the stochastic nature of the decision rule rendered simulation results more variable; to account for this we simulated 6 control networks for each of the 8 control subjects tested in the empirical study, and the same number of lesioned networks, giving 48 networks per group.

